# OPUS-Design: Designing Protein Sequence from Backbone Structure with 3DCNN and Protein Language Model

**DOI:** 10.1101/2024.08.20.608889

**Authors:** Gang Xu, Yulu Yang, Yiqiu Zhang, Qinghua Wang, Jianpeng Ma

## Abstract

Protein sequence design, also known as protein inverse folding, is a crucial task in protein engineering and design. Despite the recent advancements in this field, which have facilitated the identification of amino acid sequences based on backbone structures, achieving higher levels of accuracy in sequence recovery rates remains challenging. It this study, we introduce a two-stage protein sequence design method named OPUS-Design. Our evaluation on recently released targets from CAMEO and CASP15 shows that OPUS-Design significantly surpasses several other leading methods on both monomer and oligomer targets in terms of sequence recovery rate. Furthermore, by utilizing its finetune version OPUS-Design-ft and our previous work OPUS-Mut, we have successfully designed a thermal-tolerant double-point mutant of T4 lysozyme that demonstrates a residual enzyme activity exceeding that of the wild-type T4 by more than twofold when both are subjected to extreme heat treatment at 70°C. Importantly, this accomplishment is achieved through the experimental verification of less than 10 mutant candidates, thus significantly alleviating the burden of experimental verification process.

## Introduction

Rational protein design has emerged as a powerful tool in many applications, such as therapeutic engineering (1, 2), binder development (3–5), biosensor construction (6, 7), nanomaterial synthesis (8), and enzyme optimization (9–12). One critical sub-problem in this area is protein sequence design, also known as protein inverse folding, where the objective is to deduce the optimal amino acid sequence from a predefined backbone structure, characterized by the specific positions of backbone atoms N, C_α_, C and O. To address this problem, over the past few years, many methods have been developed.

Given the significant computational costs associated with traditional physical-based approaches, which often result in unsatisfactory accuracy (13), the research community has increasingly focused on developing deep learning models as a viable solution in the field. For example, SPIN (14) and SPIN2 (15) are two pioneering works developed by Zhou’s group, they employ several fully-connected layers as their network architectures. Meanwhile, some researchers adopt 3D Convolutional Neural Network (3DCNN) to extract distinctive features for each residue (16, 17). These methods usually necessitate a separate preprocessing step, followed by individual prediction for each residue.

Recently, numerous graph-based models have emerged and have demonstrated their potential to achieve superior sequence recovery rates in comparison to 3DCNN-based models. For example, PiFold (18) incorporates a unique residue feature extractor alongside PiGNN layers, enabling the one-shot generation of protein sequences with enhanced recovery rates. ProteinMPNN (19) and ESM-IF (20) leverage GNN-based modules to encode backbone structure and subsequently decode the corresponding sequences in an end-to-end autoregressive fashion. It is noteworthy that ESM-IF significantly augments the training data by approximately three orders of magnitude through the utilization of AlphaFold2 (21) to predict structures for 12 million protein sequences. Consequently, ESM-IF achieves a marked improvement over others. Furthermore, some researchers have incorporated protein evolutionary features derived from protein language models, such as ESM-2 (22), to further enhance the accuracy of their models (23).

It this study, we present a two-stage protein sequence design method named OPUS-Design. The first stage employs a 3DCNN-based feature extraction module, adapted from our preceding work OPUS-Rota5 (24), to capture the local environmental features of each residue. Meanwhile, the constraints from side-chain conformations are also introduced in this module. In the second stage, we use a transformer-based feature aggregation module to integrate various types of information, which include 1D protein backbone features, 2D residue contact features, and 3D local environmental features. Moreover, the protein language model, ESM-2, is incorporated for further refinement. We assess the performance of OPUS-Design against other leading sequence design methods using recently released targets from CAMEO and CASP15. The results demonstrate that OPUS-Design significantly surpasses these methods on both monomer and oligomer targets. Specifically, on the CAMEO monomer test set, which contains 288 targets, OPUS-Design achieves sequence recovery rates of 61.22% and 75.38% for all residues and core residues, respectively, compared to 55.31% and 69.54% achieved by the second-best method, ESM-IF.

Furthermore, we introduce a finetune version of OPUS-Design, designated as OPUS-Design-ft, which incorporates additional information beyond the protein backbone structure. In collaboration with our prior work, OPUS-Mut, we have successfully designed a double-point mutant of T4 lysozyme that exhibits a remarkable enhancement in residual enzyme activity, exceeding that of the wild-type T4 lysozyme by more than twofold after undergoing a 30-minute extreme heat treatment at 70°C. Notably, the design of this remarkable thermal-tolerant mutant is accomplished through the experimental validation of less than 10 candidates.

## Methods and Materials

### Framework of OPUS-Design

The architecture of OPUS-Design is methodically organized into two distinct modules, a 3DCNN-based feature extraction module and a transformer-based feature aggregation module.

In the first feature extraction module, we employ an adapted version of the 3D-Uet from OPUS-Rota5 (24) to capture the local environmental and the side-chain features of each residue. Inspired by DLPacker (25), we isolate a 20 Å box around each residue, encompassing its immediate surroundings, and subsequently align the atoms within this box to a standardized reference frame. To represent the atomic distribution within these isolated boxes, we employ a 3D Gaussian density kernel expansion, mapping each atom onto a 40×40×40 voxel grid. We use 27 channels for representation: five dedicated to elemental identities (C, N, O, S, and others), one to encode partial charge information derived from the Amber99sb force (26), and 21 to distinguish between residue types (20 channels for common residues and one for others). The outputs of the first module include three branches. The first branch, comprising four channels (C, N, O, and S), depicts the 3D side-chain density of the target residue. The second branch, with two channels, quantifies the likelihood of each voxel within the grid containing side-chain density. Lastly, the third branch, containing 20 channels, offers the probability of each of the common residue types in the box. A detailed illustration of the first module is provided in Figure 1A.

**Figure 1.**
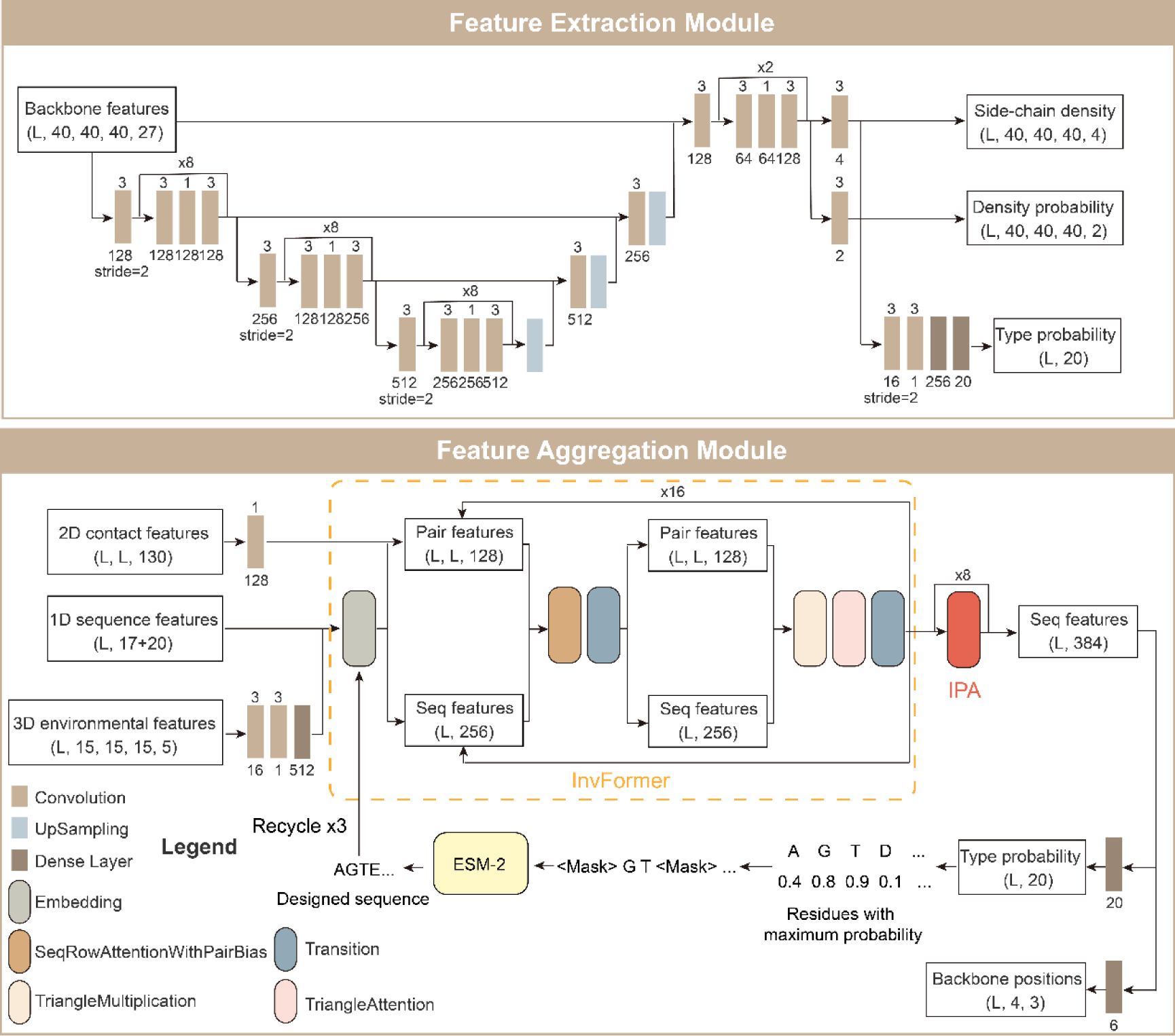
Overview of the OPUS-Design workflow. A. The first stage employs a 3D Convolutional Neural Network (3DCNN) for feature extraction, specifically designed to capture the 3D local environmental features of each residue. This module enables predictions to be made solely based on the local structural context. B. The second stage adopt a transformer-based feature aggregation module to integrate diverse feature types: 1D protein sequence features, including the predicted probability distributions of each type of residues obtained from the first stage, 2D residue-residue backbone contact features, and the aforementioned 3D local environmental features. Subsequently, 8 IPA modules are deployed to reconstruct the 3D coordinates of backbone atoms. Furthermore, the workflow incorporates a refinement step utilizing ESM-2, where positions with a maximum predicted probability below 0.5 are designated as “mask” encodings, thereby undergo further optimization.

During the training process of the first module, we adopt the mean absolute error (MAE) loss between the predicted and actual side-chain densities for the first branch, and the cross-entropy loss for the second and the third branches. Adam optimizer (27) is used for optimization with an initial learning rate of 1e-3. To mitigate overfitting problem, we implement a learning rate scheduling strategy. Specifically, if the accuracy on the validation set declines following the completion of an epoch, we halve the learning rate. Training ceases after the rate has been reduced four times. We adopt an ensemble learning approach, training five independent models using the aforementioned configuration. The final prediction is determined as the average of their outputs. Each model is trained on four NVIDIA Tesla V100s.

In the second feature aggregation module, we integrate three distinct feature sets utilizing a transformer-based InvFormer module. In addition, we incorporate the IPA module, which is derived from AlphaFold2 (21), to introduce structural constraints. The three feature sets include: 1) 1D protein sequence features, which encompass 3-state and 8-state secondary structures, alongside 6 backbone torsion angle features (consisting of sine and cosine values for ϕ, ψ, and ω angles). 2) 2D residue-residue backbone contact features, sourced from trRosetta (28), which include the C_β_-C_β_ distance distributions and the orientational distributions of three dihedrals (ω, θ_ab_, θ_ba_) and two angles (φ_ab_, φ_ba_) between residues a and b. The distance distributions span from 2 to 20 Å, segmented into 36 bins at 0.5 Å intervals, with an additional bin accounting for distances exceeding 20 Å. The φ angle ranges from 0 to 180°, segmented into 18 bins at 10° intervals, inclusive of a non-contact bin. Both ω and θ vary from −180 to 180°, segmented into 36 bins at 10° intervals, also with a non-contact bin. Here, the positions of C_β_ atoms are determined utilizing the coordinates of backbone atoms. 3) 3D local environmental features, which come from the first feature extraction module, comprising 5 channels. The first channel contains the local information of each residue, obtained by culminating the density of backbone atoms in the corresponding box. The remaining 4 channels are the predicted 3D side-chain density of target residue, obtained from the first output branch of the feature extraction module. For higher efficiency, we exclude the peripheral bins (specifically, 5 bins from each direction) and down-sample the density map from (40, 40, 40, 5) to (15, 15, 15, 5). Furthermore, the probability distribution for each residue type from the third output branch of the feature extraction module are also included.

As shown in Figure 1B, the InvFormer module is utilized to integrate both sequence and pair features. To construct the input for sequence features, the 3D local environmental features are initially processed through two 3D convolutional layers, subsequently flattened, and concatenated with 1D protein sequence features and probability distributions derived from the first feature extraction module. Concurrently, the 2D residue-residue backbone contact features undergo a 2D convolutional layer to formulate the input for pair features. In OPUS-Design, 16 InvFormer modules are deployed to enhance feature extraction capabilities. After feature aggregation, 8 IPA modules are employed to reconstruct the 3D coordinates of backbone atoms, making sequence features to encapsulate the structural information. The outputs of the feature aggregation module bifurcate into two branches: one dedicated to predicting the probabilities of 20 common residue types, and the other focused on predicting backbone conformations, utilizing the FAPE loss from AlphaFold2 as a guidance. Notably, the probability distributions for each residue type, along with the sequence and pair features from the preceding iteration, are reintegrated into the initial inputs, and this entire process is reiterated three times.

Same as the training approach of the feature extraction module, we utilize Adam optimizer with a starting learning rate at 1e-4. Furthermore, we train three distinct models and determine the final prediction by calculating the average of their respective outputs.

Additionally, we introduce the protein language model, ESM-2 (22), to deliver refined results from an evolutionary standpoint. The protein language model is integrated into both feature extraction and aggregation modules. Briefly speaking, we designate the predicted residues with low confidence as “mask” and proceed to replace them with the outputs generated by ESM-2.

### Finetune version of OPUS-Design (OPUS-Design-ft)

In this study, we introduce an enhanced version of OPUS-Design, namely OPUS-Design-ft, which undergoes a fine-tuning process. When confront with protein engineering tasks where both the amino acid sequence and experimentally determined structure of the target protein are available, a pivotal objective arises: to enhance the thermostability or enzyme activity of target protein. To address this, we have developed OPUS-Design-ft to incorporate additional information, specifically the side-chain conformations and original sequence data, beyond the exclusive reliance on backbone structure in the original OPUS-Design.

OPUS-Design-ft only retains the first 3DCNN-based feature extraction module from OPUS-Design, but with some crucial modifications: it now incorporates not only the backbone atoms but also the side chains of surrounding residues in the box, thereby integrating side-chain information into the model. Moreover, whereas OPUS-Design employs 21 channels (20 for common residues and one for others), initially setting the residue type for all atoms in the box to the “others” category, OPUS-Design-ft initializes each atom with its specific, original residue type, while only masking the type of predicted residue for training purpose. This approach ensures that the original sequence information is effectively utilized.

Consistent with the training approach employed in OPUS-Design, we adopt the Adam optimizer, initiating with a learning rate of 1e-4. Three distinct models are trained using the pre-trained models from OPUS-Design as their starting checkpoints.

### Datasets

In OPUS-Design, we employ the same training dataset as utilized by trRosetta (28), consisting of 15,051 protein targets released before May 2018. To evaluate the performance across various methods, we adopt two distinct test sets comprising monomer targets: CAMEO and CASP15. CAMEO contains 288 monomer targets, sourced from the CAMEO website (29), spanning the recent four years, in which 65 released between May 2021 and October 2021, 81 released between April 2022 and June 2022, 82 released between May 2023 and August 2023, and 60 released between March 2024 and June 2024. The CASP15 set, available for download from the CASP website (http://predictioncenter.org), comprises 44 additional monomer targets. To facilitate structural evaluation, we have constructed a test set named CAMEO-TM90, utilizing the targets sourced from the aforementioned CAMEO dataset. This test set comprises 137 monomer targets, each of which exhibits a TM-score exceeding 0.9 when comparing their native structures to the respective predicted structures obtained through ESMFold (22). Furthermore, we have gathered a test set for oligomer targets, CAMEO-oligomer, consisting of 51 oligomer targets released between March and June 2024 by the CAMEO website.

### Performance Metrics

In this study, we adopt the sequence recovery rate as the primary quantitative metric, specifically quantifying the proportion of correctly predicted residues between the designed and native protein sequences. A high recovery rate signifies a close similarity between the designed and native sequences. Furthermore, we utilize the self-consistent TM-score (sc-TM), a metric that measures the structural similarity between predicted structures generated using the native sequence and the designed sequence. For the purposes of structural evaluation, we have limited our analysis to targets that demonstrate a TM-score exceeding 0.9 when comparing the native structure to the prediction obtained from ESMFold. This threshold is essential to ensure that we are comparing against reliable structural predictions, as an elevated structural similarity towards unreliable structures may ultimately yield misleading conclusions.

In this study, in accordance with the FASPR (30), a residue is defined as a core residue when there exist more than 20 residues within a 10 Å C_β_-C_β_ distance threshold (with C_α_ considered for Glycine residues). Furthermore, we classify residues as interfacial residues if they possess at least one neighboring residue, located on a distinct peptide chain, with a C_α_–C_α_ distance of less than 8 Å.

### Data and Software Availability

The code and pre-trained models of OPUS-Design as well as the test sets used in the study can be downloaded from http://github.com/OPUS-MaLab/opus_design. They are freely available for academic usage.

### Protein expression and purification

The DNA sequences coding for bacteriophage T4 lysozyme (UniProt ID: P00720) wild-type (WT) and mutants are individually synthesized and inserted into the pET-45b (+) vector, which features an N-terminal His_6_-tag for protein purification. Following successful insertion, these recombinant vectors are transformed into BL21(DE3) chemically competent cells separately. Selected colonies are cultured in LB medium containing 100 μg/ml ampicillin at 37°C with shaking at 225 rpm until an OD600 of 0.6-0.8 is reached. Protein expression is induced with 0.5 mM isopropyl β-D-1-thiogalactopyranoside (IPTG) at 30 °C for 2.5 hours. Subsequently, cells are harvested by centrifugation at 3,500 rpm for 20 minutes.

Cell pellets are resuspended in Buffer A (30 mM HEPES pH 7.6,150 mM NaCl, 10 mM imidazole, 0.5 mM TCEP), with a protein inhibitor cocktail and a suitable amount of nuclease, and then lysed by sonication. After centrifugation, the clarified supernatant is loaded onto a gravity column packed with Ni-NTA agarose (QIAGEN). The column is washed with Buffer B (30 mM HEPES pH 7.6, 150 mM NaCl, 20 mM imidazole, 0.5 mM TCEP). Bound proteins are eluted using Buffer C (30 mM HEPES pH 7.6, 150 mM NaCl, 250 mM imidazole, 0.5 mM TCEP), concentrated, and then applied to a pre-equilibrated Superdex75 Increase 10/300 GL (GE Healthcare) with Buffer D (30 mM HEPES pH 7.6, 150 mM NaCl, 0.5 mM DTT) for further purification. Peak fractions containing the target proteins are pooled and concentrated for biochemical experiments. All protein purification steps are carried out at 4°C.

### Lysozyme activity assay

To assess lysozyme activity, the EnzChek^®^ Lysozyme Assay Kit (Thermo Fisher) is utilized. The substrate is prepared at a concentration of 50 μg/mL using the reaction buffer provided in the kit. Both WT and mutant protein samples are diluted separately in the reaction buffer supplemented with 1 mM TCEP, each to a specific concentration within the 0.4-4 µM range. Control samples consist of buffer only. Wells of a 96-well plate (Costar^®^ 3916) receive 50 µL of the prepared 50 μg/mL substrate, followed by 50 µL of the experimental samples. The mixtures are gently mixed, and the plate is then immediately placed into a microplate reader for fluorescence detection. Fluorescence release is monitored every 5 minutes using a BioTek Synergy LX with an excitation wavelength of 485±20 nm and an emission wavelength of 528±20 nm at room temperature (RT). Baseline fluorescence is established by measuring the fluorescence of the control samples and subtracting it from the readings of all experimental samples. Each sample is tested in triplicate. Additionally, the residual enzyme activity of each sample is examined. Specifically, each sample undergoes a process of heating at distinct temperatures for a duration of 30 minutes. After centrifugation, the enzymatic activity assay is performed at RT. The activity of untreated WT T4 lysozyme at RT is used as the 100% reference point. The relative activity is determined using the following equation:

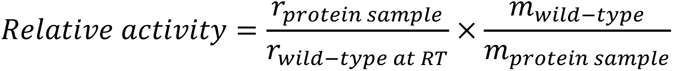

where *r* represents the slope of fluorescence increase during the initial 20 minutes, and *m* denotes the protein mass used in the assay.

### Thermal shift assay

The midpoint temperature (Tm) of the thermal denaturation curve, determined using a thermal shift assay (TSA), serves as an important indicator of the stability of protein conformation. Protein samples, either WT or mutant, are mixed with the Protein Thermal Shift™ Dye (Thermo Fisher) according to the manufacturer’s protocol, resulting in a final protein concentration of 5 µM per sample. The TSA is conducted using a QuantStudio™ 6 Flex Real-Time PCR System, with a temperature ramp from 25°C to 99°C at a rate of 0.05°C/s. The x2(520±10)-m2(558±11) optical filter is selected to capture the ROX fluorescence signal during denaturation. Each sample is tested in four replicates. The fluorescence curves obtained by truncating the post-peak region are fitted to the Boltzmann equation to determine the Tm value (31).

## Results and Discussion

### Performance of different methods on monomer targets

We evaluate the performance of OPUS-Design against several leading methods, namely ProteinMPNN (19), PiFold (18) and ESM-IF (20), utilizing two monomer test sets: CAMEO and CASP15. The results show that OPUS-Design consistently outperforms the competing methods in terms of the sequence recovery rate, both when considering all residues and focusing specifically on core residues (Table 1). It is worth noting that the training data employed by ESM-IF is two orders of magnitude larger than that utilized by the other methods.

**Table 1.** The comparison of sequence recovery rates achieved by different methods on the CAMEO and CASP15 test sets, measured by all residues and core residues.

|  | CAMEO |  | CASP15 |  |
| --- | --- | --- | --- | --- |
|  | all residues | core residues | all residues | core residues |
| ProteinMPNN | 46.39% | 57.71% | 39.51% | 53.14% |
| PiFold | 50.20% | 62.33% | 42.72% | 57.61% |
| ESM-IF | 55.31% | 69.54% | 48.28% | 66.62% |
| OPUS-Design | 61.22% | 75.38% | 50.89% | 66.81% |

In the practical realm of protein engineering, it is essential to recognize that a successful design process frequently necessitates the integration of diverse methodologies (32). Thus, while the superior performance of an individual method is undoubtedly significant, its ability to synergize with outcomes from other methods emerges as an even more crucial consideration. By integrating the results of ESM-IF and OPUS-Design, and focusing our analysis on residues that exhibit congruent predictions across both methods, we have identified that 57% of all residues and 68% of core residues share identical predictions in the CAMEO test set. For these specific residues, we have achieved an enhanced accuracy of 80.65% for all residues and 88.96% for core residues, respectively. These observations highlight the complementary strengths of OPUS-Design, establishing it as a valuable tool within the practical realm of protein engineering and design.

### Performance of different methods on oligomer targets

Additionally, we assess the performance of different methods on the oligomer test set CAMEO-oligomer. Note that, the current versions of PiFold and ESM-IF cannot directly process targets comprising multiple chains. Consequently, to accommodate these methods within our evaluation framework, we partitioned the oligomer targets into individual monomers for subsequent computational analysis. As shown in Table 2, OPUS-Design demonstrates superior performance relative to the other methods, exhibiting higher sequence recovery rates, both inclusively across all residues and specifically among interfacial residues.

**Table 2.**
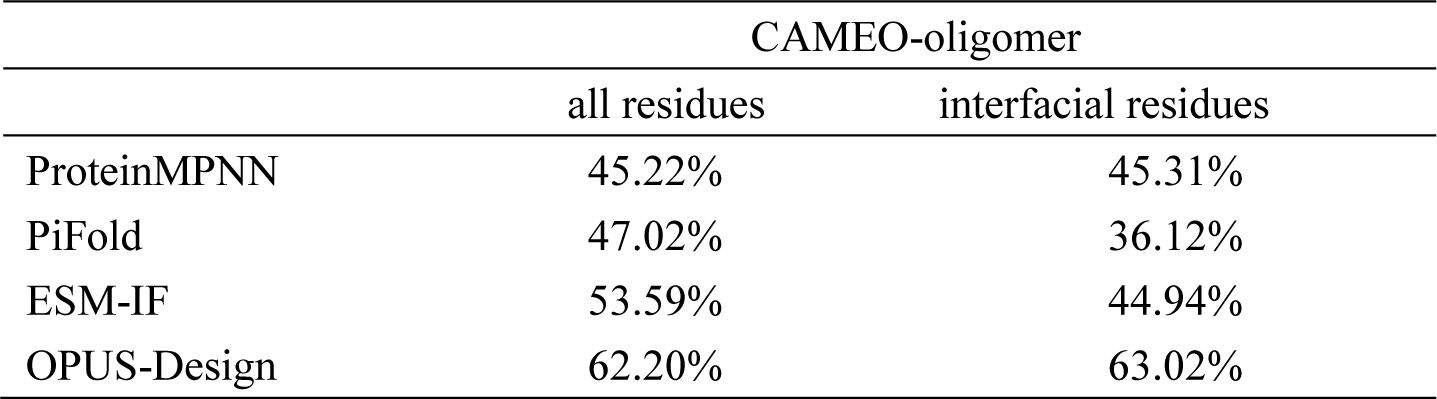
The comparison of sequence recovery rates achieved by different methods on the CAMEO-oligomer test sets, measured by all residues and interfacial residues.

Meanwhile, the decline in performance observed for PiFold and ESM-IF in predicting interfacial residues underscores the importance of incorporating local environmental information derived from neighboring chains in achieving optimal prediction accuracy.

### Ablation study of OPUS-Design

To investigate the individual contribution of each module within OPUS-Design, we conduct an ablation study by assessing the performance in the absence of specific components. OPUS-Design comprises two distinct stages: firstly, a 3DCNN-based feature extraction module, tasked with capturing the local environmental features of each residue; secondly, a transformer-based feature aggregation module, responsible for integrating diverse types of features. Furthermore, evolutionary insights are incorporated through the utilization of the protein language model, ESM-2. As shown in Figure S1, the exclusion of information from any of these components leads to a marked decline in OPUS-Design’s performance. Moreover, the results underscore the importance of the 3DCNN-based feature extraction module, as it captures the most crucial local environmental features that significantly contribute to the overall performance of OPUS-Design.

### Evaluating through structure prediction methods may be imprecise

In protein sequence design, many studies often incorporate a predicted structure comparison task to showcase the enhanced performance of their proposed methods. However, we argue that the utilization of structural indicators for assessing the efficacy of sequence design approaches is subject to imprecision due to intrinsic limitations in structure prediction methods. As shown in Table 3, we compare the structural similarity between predicted structures generated using the native sequence and the designed sequence for each method, utilizing ESMFold (22). The results show that, while the recovery rates exhibit notable variations among the different methods, the sc-TM scores for each method are very close. Given the intrinsic errors associated with structure prediction methods, the subtle differences observed in structural similarity performance may not necessarily signify meaningful distinctions in the evaluation of sequence design methods.

**Table 3.**
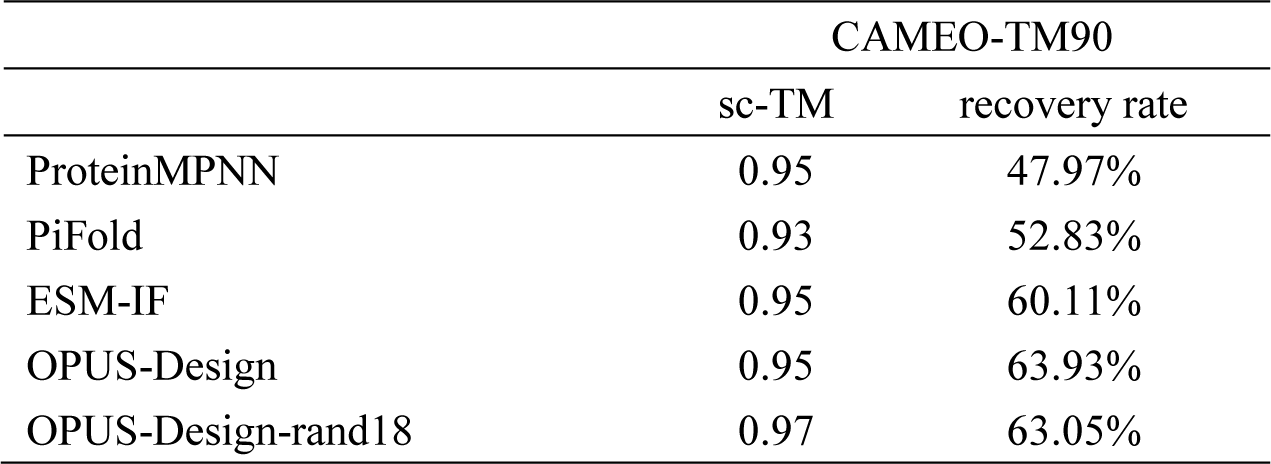
The comparison of sc-TMs and sequence recovery rates achieved by different methods on the CAMEO-TM90 test set.

Most importantly, since we have already known the target structure, we can enhance the sc-TM score by iteratively executing the program and selecting the sequence that exhibits the highest structural similarity to the target structure. To incorporate randomness into OPUS-Design, we randomly mask the local environmental features of several residues before introducing them into the feature aggregation module. By iteratively executing the second stage of OPUS-Design 18 times, we obtain 18 distinct sequences. Our findings, as presented in Table 3 (OPUS-Design-rand18), indicate that by selecting the predicted structure with the highest structural similarity to the target structure, OPUS-Design-rand18 achieves the highest sc-TM value of 0.97, albeit with a slight decrease in sequence recovery rate compared to OPUS-Design.

Bacteriophage T4 lysozyme, a monomeric protein consisting of 164 amino acid residues, possesses the enzymatic capability to hydrolyze peptidoglycan, with E11, D20, and T26 identified as the active site residues (33). Within this protein, we have selectively preserved the residues spanning from positions 10 to 30 while randomly mutating a subset of the remaining residues. Consequently, 8 mutants are constructed, with their detailed sequences listed in Table S1. As shown in Table 4, upon analysis of the structural similarity between the native T4 lysozyme structure and the 8 predicted mutant structures generated by ESMFold (22), we observe a generally high level of congruence. Specifically, the sc-TMs for all 8 mutants surpass 0.90, with 7 of these mutants achieving sc-TM exceeding 0.95. Consequently, from a structural perspective, these mutants appear to be promising candidates.

**Table 4.** The structure prediction results and the level of soluble expression for 8 mutants of T4 lysozyme. “2lzm” refers to the results of wild-type bacteriophage T4 lysozyme. “TM-score” represents the structural similarity between the experimentally determined wide-type T4 structure and the predicted structure generated using the corresponding sequence by ESMFold.

|  | # of mutations | recovery rate | TM-score | sc-TM | level of soluble expression* |
| --- | --- | --- | --- | --- | --- |
| 2lzm | 0 | 100.00% | 0.99 | 1.00 | +++ |
| 2lzm_#1 | 10 | 93.90% | 0.98 | 1.00 | +++ |
| 2lzm_#2 | 10 | 93.90% | 0.89 | 0.90 | ++ |
| 2lzm_#3 | 11 | 93.29% | 0.94 | 0.95 | + |
| 2lzm_#4 | 19 | 88.41% | 0.98 | 1.00 | + |
| 2lzm_#5 | 19 | 88.41% | 0.99 | 1.00 | + |
| 2lzm_#6 | 20 | 87.80% | 0.98 | 1.00 | - |
| 2lzm_#7 | 30 | 81.71% | 0.98 | 0.99 | - |
| 2lzm_#8 | 53 | 67.68% | 0.94 | 0.96 | - |

\* “+++” denotes high solubility, “++” denotes moderate solubility, “+” denotes low solubility, and “-” denotes nearly complete insolubility, with proteins forming inclusion bodies.

However, upon the construction of *Escherichia coli* expression vectors for these proteins and the subsequent examination of their expression levels (Figure S2 and Table 4), we observe that while 2lzm, 2lzm_#1 and 2lzm_#2 exhibit a favorable soluble expression profile, the remaining proteins either exhibit minimal soluble expression or are predominantly confined within inclusion bodies, potentially due to abnormal folding patterns. In addition, our findings indicate a correlation between the reduced recovery rates of mutant proteins and their heightened tendency to express in the form of inclusion bodies.

A study from Baker’s group show that, the absence of evolutionary constraints in the design of TEV protease variants results in a loss of enzymatic activity towards peptide substrates, emphasizing the pivotal role of conserving critical residue (34). However, in the context of protein inverse folding, where only the backbone structure is known in advance, the information regarding conserved residues remains elusive. Therefore, maximizing the sequence recovery rate in protein inverse folding may be crucial in protein design.

### Designing T4 lysozyme with OPUS-Design-ft

In this case study, we download the structural coordinates of the wild-type T4 lysozyme (PDB ID: 2LZM) (35), which includes 164 residues. We designate 7 residues as special residues, with three of them (E11, D20, and T26) being directly associated with the active site, and the remaining four (L32, F104, S117, and N132) playing a pivotal role in the binding site. Additionally, we have identified 41 surrounding residues, which are defined as those residues that have at least one neighboring special residue located within a C_β_-C_β_ distance threshold of less than 8 Å.

We utilize OPUS-Design-ft to predict the possibility of 20 residue types at each position for all 164 × 19 = 3116 potential single-site mutations. Subsequently, we sum up the probabilities of the correct residue type, as determined by the wild-type T4 sequence, for three distinct sets of residues: 7 special residues, 41 surrounding residues, and all 164 residues respectively. For each mutant, we calculate the difference of probabilities between the mutant and the wild-type T4 for three distinct residue sets. These differences are denoted as *S_diff_spe_*, *S_diff_sur_* and *S_diff_all_*, respectively, and serve as indicators for evaluating the suitability of mutations *for* selection. Specifically, larger values of these indicators correspond to higher confidence in the correct prediction. Consequently, we employ a selection criterion that prioritizes mutations exhibiting relatively large values of these indicators for further analysis.

A recent work has demonstrated the potential advantages of utilizing composite metrics derived from diverse methodologies in protein design (32), thus in this study, we incorporate the outcomes from another method OPUS-Mut (36, 37) into our analysis, which include the predict side-chain conformations and their associated prediction confidences (pRMSD) for each residue. Based on our previous work (36), we calculate the summation of differences (*S_diff_*___*_anlge_*) in side-chain dihedral angles between the predicted wild-type and mutant side chains across all residues. This metric serves as an indicator of the potential structural perturbation caused by a given mutation, with larger *S_diff_*___*_angle_* values corresponding to more severe disruptions and potentially increased likelihood of stability loss. Consequently, in this study, we employ another criterion of selecting mutations with relatively small *S_diff_*___*_angle_* values.

The predicted Root Mean Square Deviation (pRMSD) serves as a metric to quantify the confidence in OPUS-Mut’s side-chain prediction. Specifically, a residue with a lower pRMSD value indicates a higher degree of certainty in OPUS-Mut’s prediction in accordance with its local environmental context (37). Therefore, the summation of pRMSD can be utilized to evaluate the potential impact of mutating particular residues by comparing them with their native counterparts. For each mutant, we calculate the difference of the overall pRMSD between the mutant and the wild-type T4, denoted as *S_diff_pRMSD_*. Consequently, we incorporate a third selection criterion that prioritizes mutations exhibiting a lower *S_diff_pRMSD_*.

During the calculation of *S_diff_*___*_angle_* and *S_diff_pRMSD_*, we sum up the respective values across all residues, applying a multiplicative scaling factor of 6 to the values associated with special residues and a factor of 2 to those of surrounding residues. This approach ensures that the importance of special residues and their immediate neighbors are appropriately weighted in the overall summation.

In this study, we select the mutants among the entire set of 3116 potential candidates according to the following two criteria. Firstly, we select mutations that are incorrectly predicted by OPUS-Design-ft, with specific conditions that require the *S_diff_spe_*, *S_diff_sur_* and *S_diff_all_* to be greater than or equal to zero, the *S_diff_*___*_angle_* to be less than 2, and the *S_diff_pRMSD_* to be less than or equal to zero. This first criterion results in the selection of three mutants: P37S, A93P, and N163D. In the second criterion, we sort the remaining mutations that adhered to the aforementioned specific conditions based on their *S_diff_spe_*, *S_diff_sur_* and *S_diff_all_* values, in descending order. Then we select the top four mutants: K35V, S90G, G113E, and T109K. Additionally, for comparative analysis, we include two extreme cases, Y18R and K48W. While the mutant Y18R possesses a large *S_diff_*___*_angle_* value that is unfavorable to our selection criteria, it possesses high values of *S_diff_spe_*, *S_diff_sur_* and *S_diff_all_*, making it a suitable candidate for inclusion. In addition, the mutant K48W is selected based on its noteworthy attribute of exhibiting the lowest *S_diff_pRMSD_* value among those that fulfill our second criterion, despite occupying a lower ranking position. In summary, 9 mutants are experimentally examined in this study (Table S2).

We have expressed and purified the wild-type bacteriophage T4 lysozyme along with 9 mutant proteins in vitro, as depicted in Figure S3. Subsequently, we employ a thermal shift assay to determine the midpoint temperature (Tm) of thermal denaturation for each of these proteins. In addition, their enzyme activities are also being examined. As shown in Table 5, after excluding two extreme mutants that served solely for comparative purposes, three of the seven mutants (A93P, G113E, and T109K) exhibit higher Tm values compared to the wild-type T4, while another three mutants (A93P, S90G, and T109K) show enhanced enzyme activity at room temperature. For two extreme mutants, the mutant Y18R display a relatively low enzyme activity of merely 3.0% compared to the wild-type T4, which can be attributed to its substantial *S_diff_*___*_angle_* value, underscoring the significance of selecting mutants with lower *S_diff_*___*_angle_* values. Additionally, the performance of the mutant K48W, representing another extreme case, is also inferior to that of the wild-type T4.

**Table 5.**
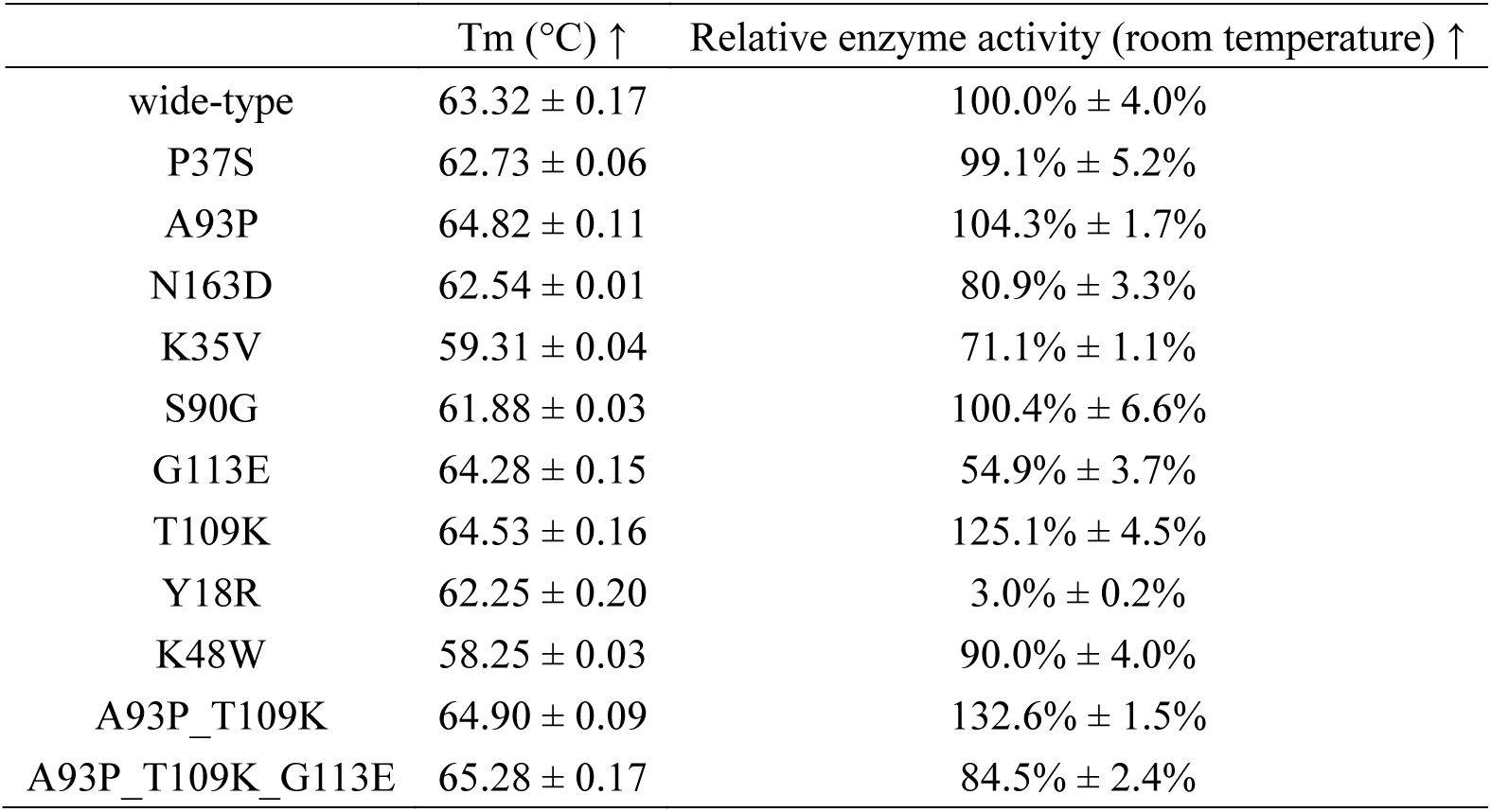
The Tm values and relative enzyme activity at room temperature for 9 selected single-site mutants, the double-point mutant A93P_T109K, and the triple-point mutant A93P_T109K_G113E of T4 lysozyme. The results are reported as mean ± SD.

| | $T_m$ ($^{\circ}\text{C}$) ↑ | Relative enzyme activity (room temperature) ↑ |
| --- | --- | --- |
| wide-type | $63.32 \pm 0.17$ | $100.0\% \pm 4.0\%$ |
| P37S | $62.73 \pm 0.06$ | $99.1\% \pm 5.2\%$ |
| A93P | $64.82 \pm 0.11$ | $104.3\% \pm 1.7\%$ |
| N163D | $62.54 \pm 0.01$ | $80.9\% \pm 3.3\%$ |
| K35V | $59.31 \pm 0.04$ | $71.1\% \pm 1.1\%$ |
| S90G | $61.88 \pm 0.03$ | $100.4\% \pm 6.6\%$ |
| G113E | $64.28 \pm 0.15$ | $54.9\% \pm 3.7\%$ |
| T109K | $64.53 \pm 0.16$ | $125.1\% \pm 4.5\%$ |
| Y18R | $62.25 \pm 0.20$ | $3.0\% \pm 0.2\%$ |
| K48W | $58.25 \pm 0.03$ | $90.0\% \pm 4.0\%$ |
| A93P_T109K | $64.90 \pm 0.09$ | $132.6\% \pm 1.5\%$ |
| A93P_T109K_G113E | $65.28 \pm 0.17$ | $84.5\% \pm 2.4\%$ |

Subsequently, we construct and purify a double-point mutant (A93P_T109K) and a triple-point mutant (A93P_T109K_G113E) for further analysis (Figure S3). As shown in Table 5, the double-point mutant A93P_T109K displays superior performance by achieving an enzyme activity that surpasses that of the wild-type T4 by 32.6% under room temperature conditions. Conversely, although the triple-point mutant A93P_T109K_G113E achieves the highest Tm among all mutants, indicative of a remarkable degree of stability, its enzyme activity is notably reduced compared to the double-point mutant A93P_T109K, falling even below that of the wild-type T4 enzyme. The observation of stability-activity trade-off phenomenon has also been reported by previous study, which is a general limitation in the design of enzymes. Specifically, enhancements in one attribute are frequently paralleled by diminutions in the other (38).

Furthermore, we conduct an evaluation of the residual enzyme activity of each mutant after subjecting them to a range of extreme temperatures (50°C, 60°C, and 70°C) for 30 minutes. As illustrated in Figure 2, the double-point mutant A93P_T109K demonstrates remarkable resilience, exhibiting a residual enzyme activity that more than twofold that of the wild-type after treatment at 70°C for 30 minutes. Notably, even after undergoing the extreme treatment at 70°C, the enzyme activity of A93P_T109K is even 17.3% higher than that of the wild-type T4, although the latter is not subjected to high thermal treatment, indicating its exceptional thermal tolerance.

**Figure 2.**
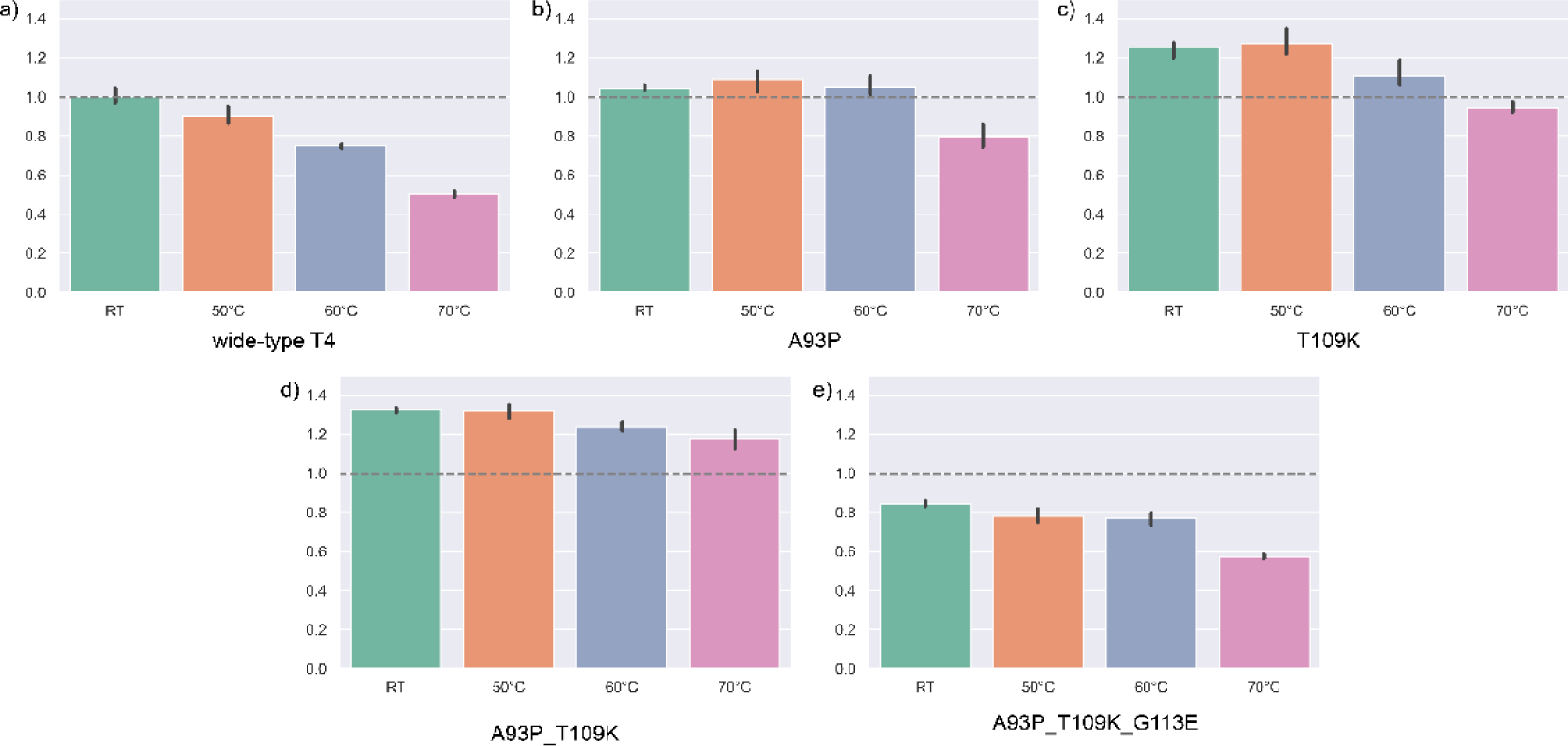
The relative residual enzyme activity of each mutant subsequent to various heat treatment conditions. The enzyme activity of the untreated wild-type bacteriophage T4 lysozyme, measured at room temperature, serves as the normalized baseline (set at 1.0), which is represented by a gray dotted line within each panel. Panels a-e) show the relative residual enzyme activities for the wide-type T4, the single-point mutant A93P, the single-point mutant T109K, the double-point mutant A93P_T109K, and the triple-point mutant A93P_T109K_G113E, respectively.

## Conclusion

In this study, we present OPUS-Design, a two-stage protein sequence design method. By integrating local environmental features captured through a 3DCNN-based module and the evolutionary features derived from ESM-2, OPUS-Design demonstrates a superiority over several leading methods in terms of sequence recovery rates across both monomer and oligomer targets. This establishes OPUS-Design as a potent instrument for applications within the realm of protein design

Furthermore, we develop a finetune version of OPUS-Design, titled OPUS-Design-ft, which incorporates additional information that transcends the backbone structure. Specifically, OPUS-Design-ft integrates side-chain conformations and original sequence data, thereby enhancing its capability to more precisely address protein engineering tasks for targets with experimentally determined structures.

It is noteworthy that enhancing enzyme activity under extreme conditions, such as high temperatures, constitutes a pivotal yet formidable challenge in protein engineering, frequently hindered by the intricate balance between stability and activity, also known as the stability-activity trade-off. Assisted by our previous work, OPUS-Mut, we have designed a double-point mutant of T4 lysozyme using OPUS-Design-ft. Our experimental results show that, subsequent to a 30-minute exposure to extreme heat treatment at 70°C, the residual enzyme activity of this mutant surpasses that of the wild-type T4 by more than twofold, indicating the promising potential of our methods in designing enzymes for high-temperature enzymatic applications. Remarkably, this mutant with exceptional thermal tolerance has been found through the experimental verification of less than 10 candidates, thereby significantly alleviating the considerable burden associated with the experimental verification process.

## Acknowledgements

J. M. wants to thank the supports from Science and Technology Innovation Plan of Shanghai Science and Technology Commission (No. 23JS1400200). G. X. wants to thank the support from National Natural Science Foundation of China (No. 32300535).

